# Different clones for different contexts: Hippocampal cognitive maps as higher-order graphs of a cloned HMM

**DOI:** 10.1101/745950

**Authors:** Nishad Gothoskar, J. Swaroop Guntupalli, Rajeev V. Rikhye, Miguel Lázaro-Gredilla, Dileep George

## Abstract

Hippocampus encodes cognitive maps that support episodic memories, navigation, and planning. Understanding the commonality among those maps as well as how those maps are structured, learned from experience, and used for inference and planning is an interesting but unsolved problem. We propose higher-order graphs as the general principle and present, as a plausible model, a cloned hidden Markov model (HMM) that can learn these graphs efficiently from experienced sequences. In our experiments, we use the cloned HMM for learning spatial and abstract representations. We show that inference and planning in the learned CHMM encapsulates many of the key properties of hippocampal cells observed in rodents and humans. Cloned HMM thus provides a new framework for understanding hippocampal function.

## Introduction

A dominant theory of information processing in hippocampus is that it encodes a cognitive map (Keefe & Nadel, 1978) that enables flexible navigation, planning and episodic memory (Tolman, 1949). A computational theory for cognitive maps should explain how context and location specific representations emerge from aliased sensory data and how the representational structure enables efficient and flexible planning. However, such a theory remains elusive. Previous attempts include modeling hippocampus as a memory index, a relational memory space, a rapid event memorizer, and systems-level models of pattern-separation and pattern-completion, but these models have not reconciled the diverse functional attributes (Moser, Kropff, & Moser, 2008; Eichenbaum, Dudchenko, Wood, Shapiro, & Tanila, 1999; Schapiro, Turk-Browne, Norman, & Botvinick, 2016) of the hippocampus under a common framework.

A recent model based on the concept of successor representation (SR) attempted to capture representational properties of place cells and grid cells (Stachenfeld, Botvinick, & Gershman, 2017) but it fails to explain several experimental observations such as the discovery of place cells that encode routes (Frank, Brown, & Wilson, 2000; Grieves, Wood, & Dudchenko, 2016), remapping in place cells (Colgin, Moser, & Moser, 2008), a recent observation of place cells that do not encode goal value (Duvelle et al., 2019), and flexible planning after learning the environment.

Here, we propose that learning higher-order graphs of sequential events with a specific representational structure might be an underlying principle that allows for flexible learning and navigation of cognitive maps. This representational structure is such that different clones of a particular receptive field differentiate themselves through lateral connections to represent specific contexts. Recently we showed that this representational structure can be instantiated as a cloned hidden Markov model (CHMM) (Dedieu et al., 2019), and that an online expectation-maximization (EM) algorithm can be used to learn higher-order sequences. Once learned, our model supports retrieval of learned sequences from partial noisy cues, goal-directed planning, and explains observed representations encoded by cells in hippocampus. In this paper, we briefly present our model, its learning, and experimental evidence in support of CHMM as a plausible model for information processing in hippocampus.

## CHMM

Cloned Hidden Markov Model (CHMM) is a generative probabilistic sequence model. It is a sparse restriction of a Hidden Markov Model (HMM) that enforces that many hidden states map deterministically to the same emission state (Fig 1A). We refer to the set of hidden states that map to the same emission state as *clones* of that emission. Dedieu et al. (2019) have shown that this sparsity enables the model to learn variable-order sequences efficiently. The probability of an observation sequence in a CHMM is

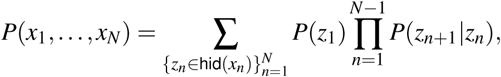

where *z*_*n*_ ∈ hid(*x*_*n*_) means that the summation is only over the hidden values of *z*_*n*_ that emit *x*_*n*_. The parameters of this model are the prior probabilities, collected in the vector π, such that π_*u*_ = *P*(*z*_1_ = *u*), and the transition probabilities *T*, such that *T*_*uv*_ = *P*(*z*_*n+1*_ = *v*|*z*_*n*_ = *u*).

**Figure 1:**
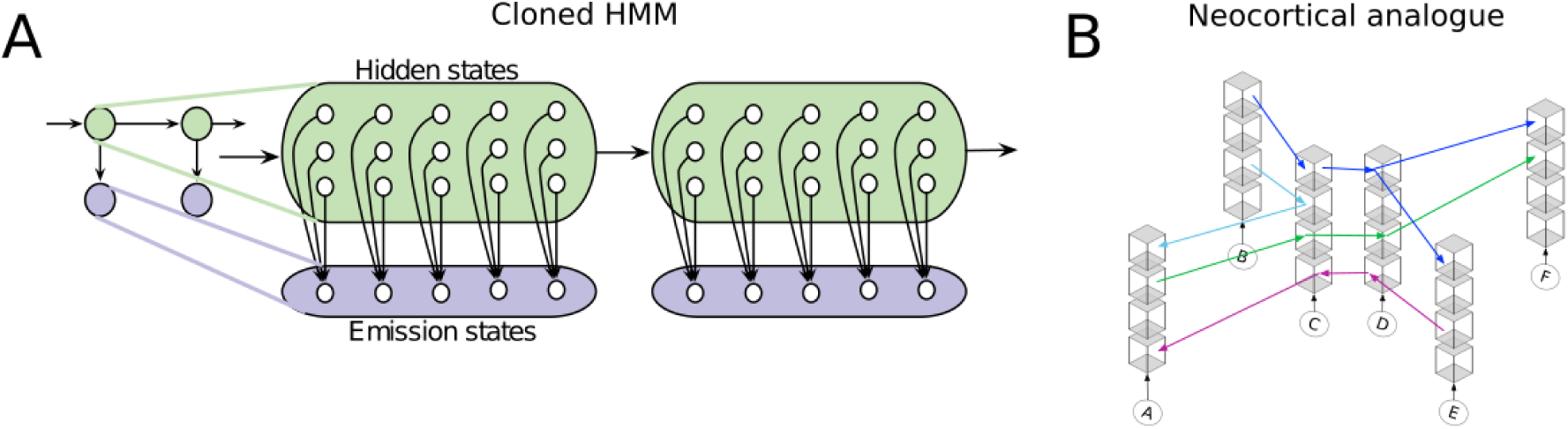
Schematic of CHMM with neural substrate. (A) Illustration of a CHMM. (B) Cortical analogy of a CHMM. Colored arrows illustrate how the cortex would represent sequences ACDF, EDCA, BCA, BCD(E,F). See text for details.

CHMMs are trained using the expectation-maximization (EM) algorithm, a coordinate ascent method for finding model parameters that maximize the likelihood of observed data. The EM algorithm alternates between computing a posterior over the unobserved variables and updating the model parameters to maximize the likelihood given those posteriors. Inference in the CHMM is exact and posteriors can be computed using forward-backward message passing. The sparse emission structure of CHMMs reduces the local minima problems during convergence and makes the messages sparse, resulting in substantially faster inference and learning, compared to HMMs (Dedieu et al., 2019).

State cloning is a mechanism for representing higher-order dependencies in a sparse manner, and the basic idea has been discovered in various domains (Hawkins, George, & Niemasik, 2009; Xu, Wickramarathne, & Chawla, 2016; Cui, Ahmad, & Hawkins, 2016; Cormack & Horspool, 1987). With a cloned representation, the same bottom-up sensory input is represented by a multitude of neurons that are copies of each other with respect to their selectivity for the sensory input. These neurons then learn to differentially activate for different temporal contexts. Fig 1B illustrates this idea for sensory inputs corresponding to locations A, B,…, F. Each column shows neurons that will respond identically to the sensory stimulus from that location in the absence of any temporal context. However, different clones are part of different temporal sequences. This kind of representation allows for the storage of a large number of higher-order and probabilistic sequences without destructive interference, enables bridging between disjoint episodes of experience, and allows for the retrieval of related sequences in an efficient manner. These properties make them suitable for representing hippocampal sequences.

## Experiments

We focus on spatial representations in our experiments but the interpretations apply to representation of abstract event structures and can potentially be extended to general sequences of events. We ran a series of experiments using CHMM for learning spatial representation using exploration in simulated environments and inference in the learned model to probe the representations of the model.

### Experiment 1: Spatial representation

In this experiment, we simulate an agent walking in the the maze shown in Fig 2A, accumulating a sequence of observations denoted by the values in each cell. The agent repeatedly walks along paths from 0 or 1 to 11. When starting at observation 0, the rat will take the upper route (passing through observation 2) and when starting from observation 1, the agent will take the lower route (passing through observation 3). A CHMM model (with 4 clones per observation) is learned from this sensory information.

**Figure 2:**
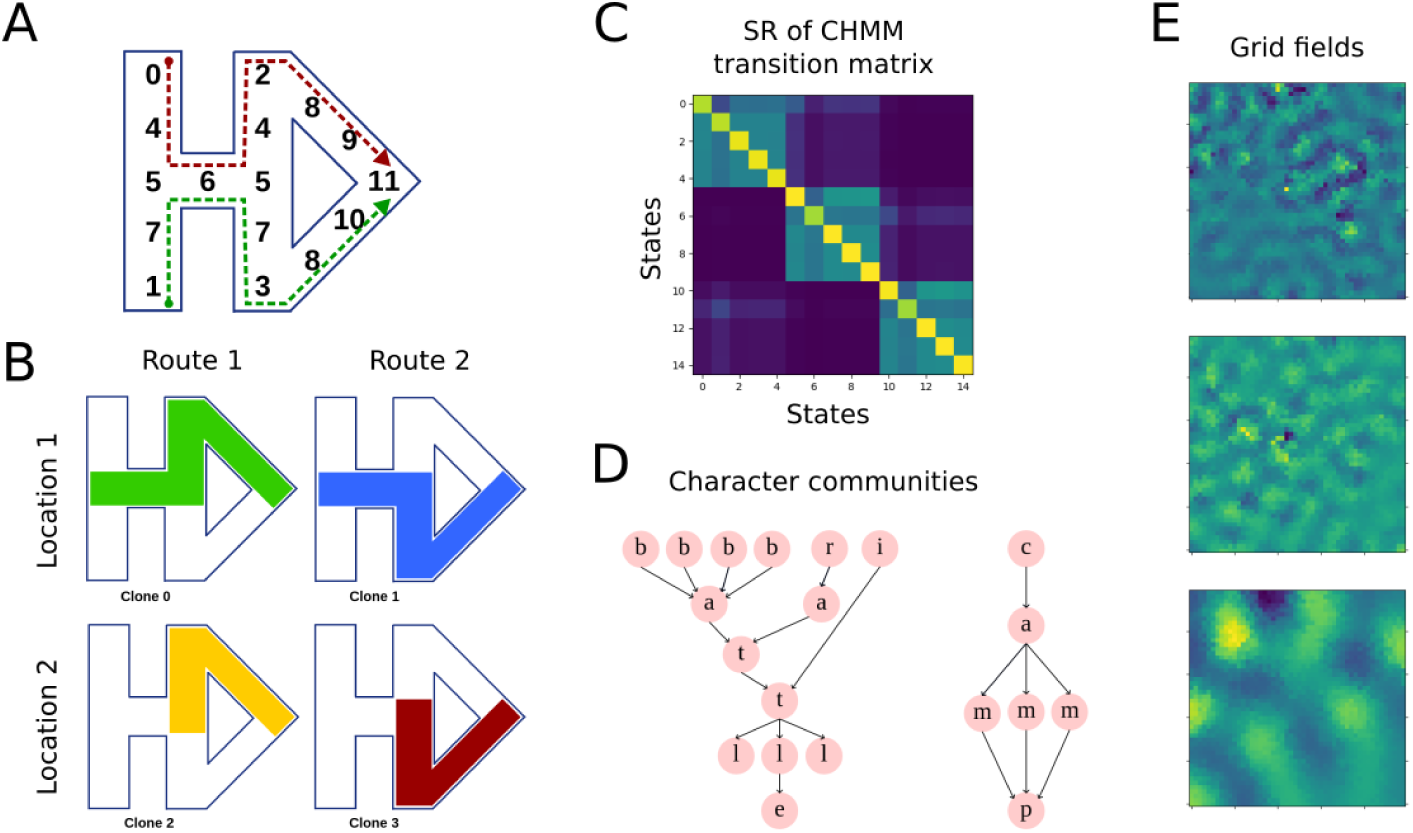
Experimental results (A) Maze used in Experiment 1 with routes experienced by the agent. Numbers indicate the sensory input at the location. (B) The clones corresponding to observation 5 encode both the different routes and locations. (C) Graph structure discovered from learned CHMM. Transition matrix reflects the graph used to generate sequences (left). (D) Character communities discovered in CHMM model trained on text (E) Eigenvectors from the CHMM transition matrix exhibit grid-like structure.

#### Route-coding

After learning the model, the agent again moves along the two routes and accumulates the corresponding observations. When running MAP inference on these observation sequences with our learned model, we observe that for each observation that is shared between the two paths (5,6,5,8) a different clone will activate depending on the route taken. This is because the clones are used to distinguish whether the shared segment will proceed into the upper or lower route. We can observe this directly by conditioning on the initial observation, 0 or 1, and sampling from the CHMM. The same effect is achieved if we condition only on 2 or 3. The produced sequence consistently correspond to the upper or lower paths, respectively. Computing the hidden state posteriors allowed us to inspect the distribution over clones for each location. For each observation that is shared between the two routes, we observed that the clone distribution for each of two were completely disjoint, showing that the clones were unique to the specific route. Together, these results encapsulate route representation (Eichenbaum et al., 1999; Frank et al., 2000; Grieves et al., 2016), and prospective coding (Grieves et al., 2016) observed in place cells.

#### State-aliasing

CHMMs can also be used for learning in settings of ambiguous sensory information. In the case of this maze, observations 4, 5, 7, and 8 appear in multiple locations and cannot by themselves dictate the agent’s physical location. Just as clones can be used to distinguish routes, they can also be used to distinguish aliased observations. Again, using the hidden state posteriors, we can inspect the clone distributions at each of the two locations. For all of these aliased observations, the distribution at the different locations were completely disjoint, showing that the clones were unique to the specific location.

Observation 5 is distinct in that it appears in two physical locations and both those locations are on the segment shared between the two routes. This requires the CHMM to use its clones to be able to both distinguish routes as well as distinguish aliased locations. Fig 2B shows 4 paths generated when conditioning on each of the clones of observation 5 and sampling forward. The CHMM properly learns to use all 4 of the clones to model both possible routes from both locations.

When a rat is dropped in a maze, it will initially be uncertain of its location within the maze. However after exploration, it will eventually be able to localize itself. In a similar fashion as described above, at first we may have a uniform distribution over clones corresponding to our initial observation, implying that we are unsure of which of the “aliased” locations we are at. After a few steps of exploration this clone distribution will collapse onto a single clone at which point we will be certain about our location. Then as the rat proceeds to explore the maze, inference at each step will allow the rat to know the clone which their current observation corresponds to.

#### Planning as inference

The learned representation of the CHMM can also be used for planning. The above statements describe that inference in the CHMM can allow the rat to know its current clone. Given the clone corresponding a goal location, finding a route from the current location to the goal corresponds to an inference problem that can be solved efficiently using message-passing algorithms (Kansky et al., 2017; George et al., 2017) or graph search.

#### Remapping

A sudden remapping of place fields and firing rates is one of the most intriguing findings about place cells (Colgin et al., 2008). Gradually changing the environment from one shape to another leads to a sudden change in place fields during intermediate morphed shapes. Such global remapping has also been observed when a salient cue in the environment is changed. These observations have been attributed to context encoding in hippocampus using attractor dynamics (Colgin et al., 2008). Remapping is implicated in the ability to encode context-dependent memory and in our model, MAP inference naturally leads to such sudden “remapping” of responses.

### Experiment 2: Graph structure

While the CHMM allows for modeling higher-order dependence, it also retains many of the useful features of first-order models like the SR. One such feature is the ability to discover community structure. To demonstrate this, we followed the experiment in Schapiro et al. (2016) in which subjects were presented with a sequence of fractals with the transition structure drawn from a graph with community structure. The subjects’ hippocampal responses showed a representational similarity with this graph, which was interpreted as hippocampus representing the event structure. We presented our model with a sequence of symbols with transitions drawn from the same graph. After learning, the SR of CHMM transition matrix (Fig 2C) showed representational similarity with the original graph. The multidimensional scaling (MDS) of dissimilarity between rows showed that clones of observations that appeared in the same community of the graph appeared in the same cluster of the MDS. A more challenging problem is to discover the higher-order structure in language. We trained a CHMM model on a character sequence from a text corpus and performed community discovery on the learned model. CHMM successfully recovered communities that corresponded to words or portions of words (Fig 2D). These results show that CHMM captures sequence learning properties of hippocampus and can efficiently learn higher-order graphs from experienced sequences.

### Experiment 3: Grid representation

Experiments have shown that a grid-like representation can be recovered from a low-dimensional embedding of place-like representation and these representations could be useful for sub-goal discovery during planning (Dordek, Soudry, Meir, & Derdikman, 2016; Moser et al., 2008). In this experiment, we performed an eigendecomposition of the CHMM representation learned from random walk in a square maze and observed that the eigenvectors exhibit a grid-like structure (Fig 2E).

## Discussion

We propose that learning higher-order graphs from the experienced events might be an underlying principle of information processing in hippocampus. We presented our recently proposed model, a cloned HMM, that can efficiently learn higherorder sequences (Dedieu et al., 2019) as a plausible model for learning, inference, and planning in hippocampus. We performed experiments simulating an agent moving in spatial environments or experiencing structured event sequences and showed how this model encapsulates observed properties of representations in hippocampus such as route-coding, remapping, encoding event transition structures, and recovering a grid-like embedding. We also showed how our learned model supports flexible planning.

Our model aligns closely with the memory space model proposed by Eichenbaum et al. (1999) in terms of a cognitive map that is more general than a spatial map. It also supports the idea of reusing events to represent different experiences that share these events albeit in different contexts (nodal coding in (Eichenbaum et al., 1999)). It supports the idea of prediction as a function of hippocampal processing (Stachenfeld et al., 2017) by being able to activate multiple likely future paths given observations (sampling possible sequences). It supports the idea of pattern-completion by being able to recover a sequence given some noisy observations. Our model subsumes some of the desirable properties of successor representation as shown by our experiments. Additionally, the CHMM can encode higher-order sequences with efficient learning that explains representations observed in hippocampus, which SR fails to do.

In this paper, we did not delve into systems-level implementation details. One future direction is to map out the specific elements and computations of our model to known hippocampal physiology and circuitry. We used a single simplistic sensory input as observations for collections of hidden states (clones), but in reality a conjunction of multiple uncertain sensory inputs feed into hippocampus. The propagation of inference messages from one sensory input to another through the clones should result in different final inferred states. This is similar to how grid cell responses change in response to sensory information being propagated through place cells. Another extension is to include multiple layers in our model. The untangled hidden states (place cells) in one level can in turn be used as inputs to another layer of hidden states, hierarchically. Inference on this network would decode not just one, but multiple levels of clones. Future experiments could verify these through simulation.

